# sideRETRO: a pipeline for identifying somatic and dimorphic insertions of processed pseudogenes or retrocopies

**DOI:** 10.1101/2020.03.09.983858

**Authors:** Thiago L A Miller, Fernanda Orpinelli, José Leonel L Buzzo, Pedro A F Galante

**Affiliations:** Centro de Oncologia Molecular, Hospital Sírio-Libanês, São Paulo 01308-060, Brazil; Departamento de Bioquímica, Universidade de São Paulo, São Paulo, Brazil

**Keywords:** Processed pseudogenes, Retrocopies, Mobile Elements, Bioinformatics, Genomics

## Abstract

Retrocopies or processed pseudogenes are gene copies resulting from mRNA retrotransposition. These gene duplicates can be fixed, somatically inserted or dimorphic in the genome. However, knowledge regarding unfixed retrocopies (retroCNVs) is still limited, and the development of computational tools for effectively identifying and genotyping them is an urgent need. Here, we present sideRETRO, a pipeline dedicated not only to detecting retroCNVs in whole-genome or whole-exome sequencing data but also to revealing their insertion sites, zygosity, and genomic context and classifying them as somatic or dimorphic events. We show that sideRETRO can identify novel retroCNVs and genotype them (93.2% accuracy), in addition to identifying dimorphic retroCNVs in whole-genome and whole-exome data. Therefore, sideRETRO fills a gap in the literature and presents an efficient and straightforward algorithm to accelerate the study of retroCNVs.

**Availability:** sideRETRO is available at https://github.com/galantelab/sideRETRO

## 1 Introduction

Studies of genetic variability caused by the insertion of long (LINE) and short (SINE) interspersed mobile elements (MEs) have been increasingly recognized as important not only for the study of evolution (Kazazian, 2004) but also that of pathologies (Burns,2017; Hancks and Kazazian, 2016). In addition to the insertion of MEs, it has been reported that cellular mRNAs are retrocopied as a byproduct of LINE retrotransposition. Humans, other primates, and mice exhibit a similar number of fixed (Navarro and Galante, 2013, 2015; Zhang *et al*., 2004) and an unclear number of unfixed retrocopies(Schrider *et al*., 2013; Ewing *et al*., 2013; Zhang *et al*., 2017; Abyzov *et al*., 2013). While the former have been well studied (Navarro and Galante, 2013; Kabza *et al*., 2014), the latter (usually referred to as retroCNVs (Schrider *et al*., 2013)) are still underexploited, especially because of the lack of a well-established algorithm for their identification.

Here, we present sideRETRO, a pipeline dedicated to detecting retroCNVs. sideRETRO uses whole-genome or whole-exome sequencing data to identify somatic or dimorphic retroCNVs and provides their genomic insertion sites, zygosity, genomic context, and parental genes.

## 2 sideRETRO: Description, main features, and availability

SideRETRO detects somatic (*de novo*) and dimorphic insertions of retrocopies that are absent in the reference genome (referred to as retroCNVs herein). SideRETRO is written in the C programming language and distributed under the GNU General Public License. It is easy to install via the command line and is also available as a Docker image (see online documentation for details).

SideRETRO is straightforward to use. It requires only an aligned BAM file (whole-genome (WGS) or whole-exome (WXS)), a reference for the genome and a reference for the transcriptome (Figure 1). For each detected retroCNV, SideRETRO provides its i) parental gene: the gene that underwent the retrotransposition process; ii) putative genomic insertion site: the genome coordinates at which retrocopy integration occurred (chromosome: start-end); iii) strandedness: the orientation of retroCNV insertion, on the same (+) or opposite (-) DNA strand compared with the parental gene; iv) genotype: when multiple genomes (individuals) are analyzed, we annotate the events occurring in each one; and v) haplotype: whether the event occurred on one (heterozygous) or both (homozygous) homologous chromosomes.

**Figure 1.**
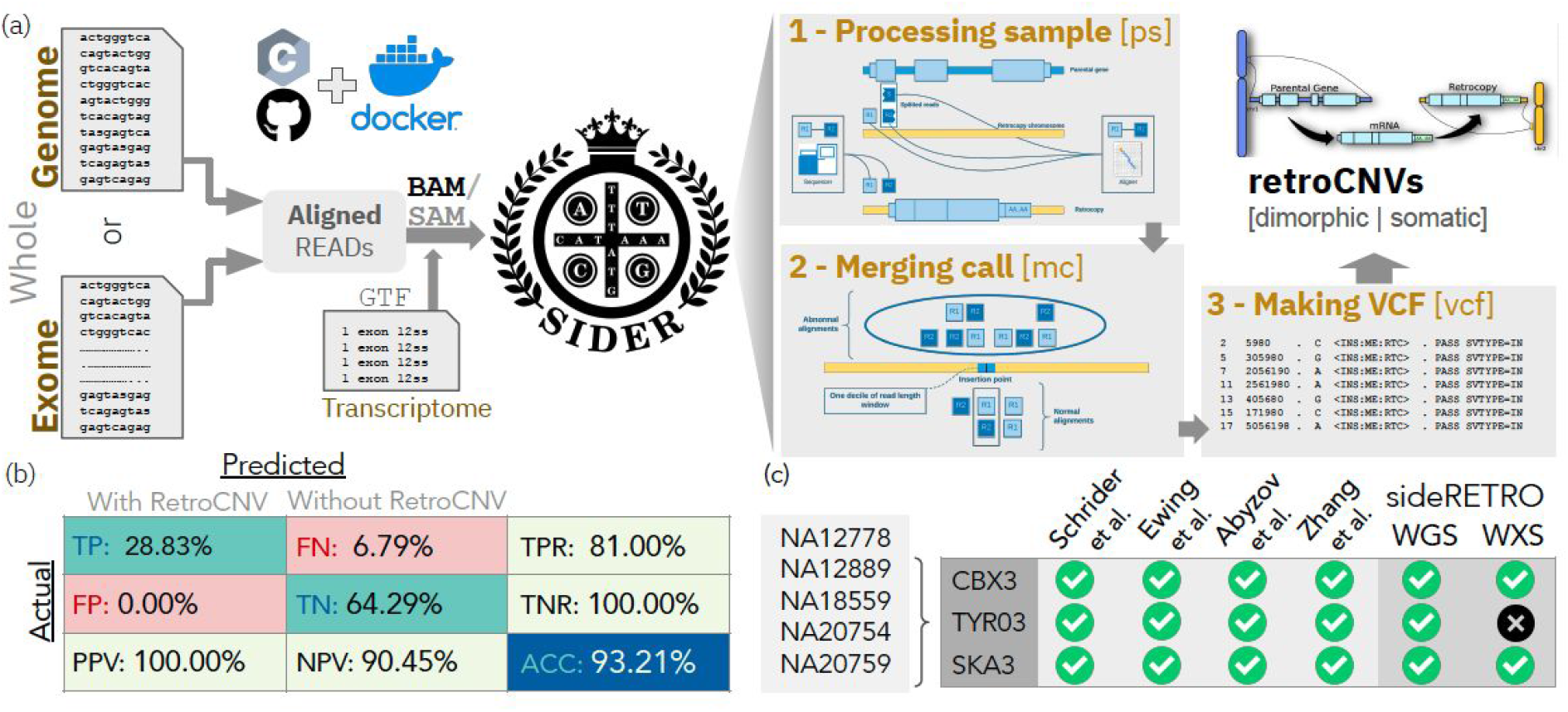
The SideRETRO pipeline is straightforward to use and accurately identifies retroCNVs. A) SideRETRO requires as input only a file with aligned reads (or list of fines) and a transcriptome annotation and then performs three major steps to identify retroCNVs and provide their characteristics (B). Confusion matrix showing the performance of sideRETRO for the simulated data. D) Consensus list of retroCNVs available in the literature(Schrider *et al*., 2013; Abyzov *et al*., 2013; Ewing *et al*., 2013; Zhang *et al*., 2017) for the individuals used here (NA12778, NA12878, NA18559, NA20754, NA29759). Events identified by sideRETRO using WGS and WXS data are highlighted in green. Abbreviations: TP: True Positive; FP: False Positive; TN: True Negative; FN: False Negative; TPR: True Positive Rate or Sensitivity; TNR: True Negative Rate or Specificity; PPV: Positive Predictive Value or Precision; NPV: Negative Predictive Value; ACC: Accuracy

SideRETRO has three subcommands: process-sample, merge-call, and make-vcf (Figure 1A-B). The process-sample subcommand reads a BAM (or SAM) file and captures abnormal reads that must be related to a retrocopy event. All of these data are saved into a SQLite3 database. In the second step, merge-call, the database is processed to detect and annotate all putative retroCNVs. Finally, the make-vcf step joins information about the identified retroCNVs and produces the sideRETRO output in VCF format (Figure 1A-B) (see online documentation for further details, https://sideretro.readthedocs.io/)).

## 3 SideRETRO: Application

To demonstrate how sideRETRO works, we used simulated datasets. We generated five human whole-genome sequencing datasets (40x coverage; (Miller et al., 2019)) with 10 randomly distributed retroCNVs in each. In total, we used a list of 30 retroCNVs (Supplementary Table 1). On average, each simulated retroCNV had at least two exons from the parental gene and was ~1000 nt in length, which are characteristics that mimic a real retrocopy (Navarro and Galante, 2015).

We applied sideRETRO to these simulated data and successfully identified retroCNVs with an accuracy of 93.2% (Figure 1C and Supplementary Figure 1). Next, we used sideRETRO to search for retroCNVs in five individuals from 1000 Genomes Project Phase 3 (1000 Genomes Project Consortium *et al*., 2015). In the WGS data, we identified 20 retroCNVs in total (Supplementary Table 2). As expected, in the WXS data from the same individuals, we identified fewer candidates: 6 retroCNVs (Supplementary Table 2). We also matched our findings to retroCNVs available in the literature (Schrider *et al*., 2013; Abyzov *et al*., 2013; Ewing *et al*., 2013; Zhang *et al*., 2017), which are three consensus events for these five individuals. SideRETRO successfully identified all of the retroCNVs with WGS data and missed only one candidate when exome data were used (Figure 1D).

## 4 Discussion

SideRETRO is a method dedicated to identifying retroCNVs in WGS or WXS data, which provides several characteristics of these gene duplicates that are not available in the literature.

We (Schrider *et al*., 2013) and others (Ewing *et al*., 2013; Abyzov *et al*., 2013; Zhang *et al*., 2017) have already searched for retroCNVs and made available the developed approaches for identifying them. However, these studies were not devoted to describing methodologies, and their pipelines were therefore limited not only in terms of their documentation, installation and use but also in terms of providing additional characteristics of retroCNV events, such as their genomic insertion sites and zygosity, and they did not present retroCNVs in a standard format such as VCF format. Moreover, sideRETRO provides key information about retroCNV events such as the parental genes, genomic insertion sites, zygosity, and genomic context (within or near a gene), and it helps to classify events as somatic or dimorphic.

In summary, we expect that sideRETRO will shed light on and drive further studies of retroCNVs in health and disease genomes.

