## Supplementary data for "sideRETRO: a pipeline for identifying somatic and dimorphic insertions of processed pseudogenes or retrocopies"

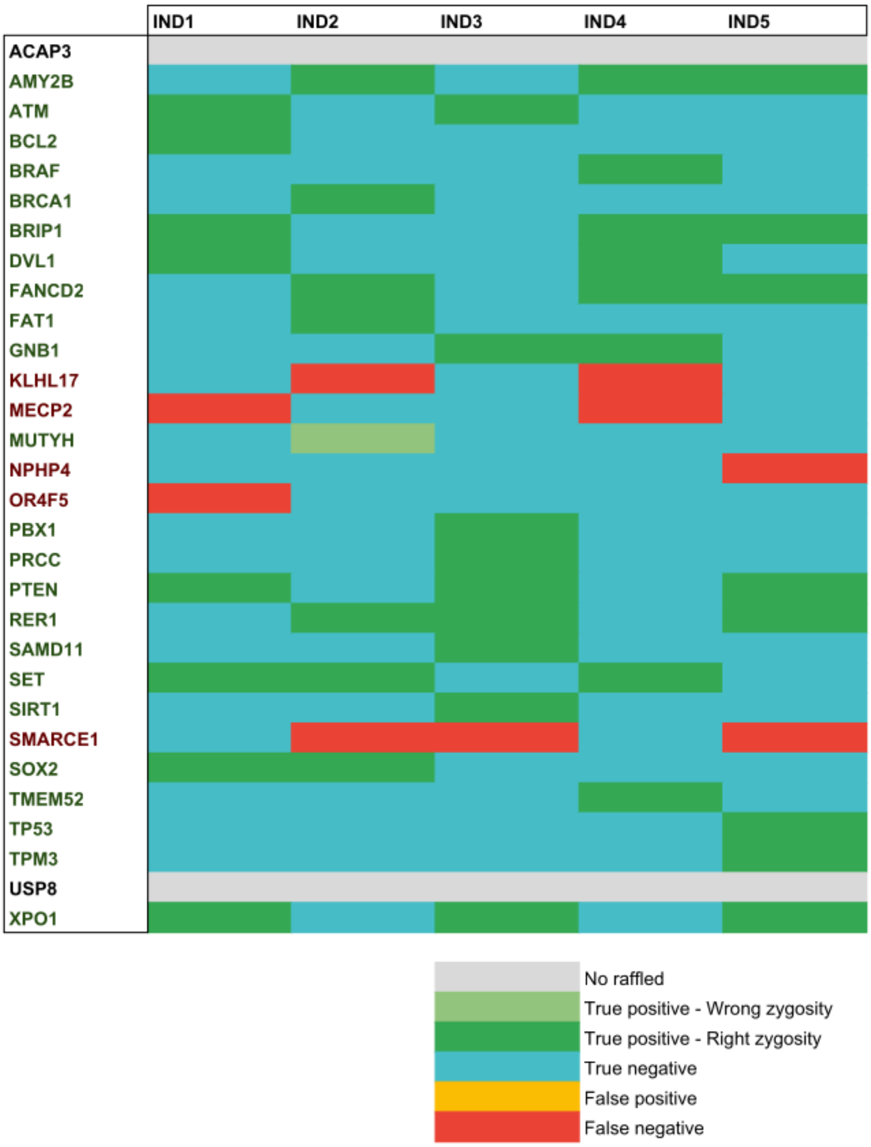


**Supplementary Figure 1. Heatmap showing the sideRETRO search results for retroCNVs in the simulated data**. Gray retroCNVs did not present reads in the simulation. IND1 to IND5 are the IDs for the five simulated genomes that we generated. We considered true-positive candidates to be those identified by sideRETRO at the same (or near +/- 10 nt) insertion site generated in the simulated data.

**Supplementary Table 1. Description of simulated and found retroCNVs.** Parental genes, positions, orientations (strand) from simulated (actual position) and found retroCNVs by sideRETRO (predicted position).

| Parental gene | Actual position | | | Predicted position | | |
|---|---|---|---|---|---|---|
| AMY2B | chr5 | 122895832 | - | chr5 | 122895831 | - |
| ATM | chr19 | 16443740 | + | chr19 | 16443738 | + |
| BCL2 | chr4 | 146453786 | + | chr4 | 146453783 | + |
| BRAF | chr18 | 22375812 | - | chr18 | 22375811 | - |
| BRCA1 | chr7 | 102911369 | - | chr7 | 102911368 | - |
| BRIP1 | chr12 | 113357216 | - | chr12 | 113357216 | - |
| DVL1 | chr5 | 54105196 | - | chr5 | 54105195 | - |
| FANCD2 | chr3 | 121339674 | + | chr3 | 121339673 | + |
| FAT1 | chr16 | 65659952 | - | chr16 | 65659951 | - |
| GNB1 | chr12 | 70734022 | + | chr12 | 70734022 | + |
| MUTYH | chr4 | 2716745 | + | chr4 | 2716760 | + |
| PBX1 | chr21 | 22718521 | - | chr21 | 22718520 | - |
| PRCC | chr1 | 190777903 | - | chr1 | 190777902 | - |
| PTEN | chr2 | 140252560 | - | chr2 | 140252559 | - |
| RER1 | chr13 | 55179109 | + | chr13 | 55179108 | + |
| SAMD11 | chr4 | 14585135 | + | chr4 | 14585134 | + |
| SET | chr7 | 154178578 | - | chr7 | 154178578 | - |
| SIRT1 | chrX | 120716688 | - | chrX | 120716687 | - |
| SOX2 | chr10 | 88689163 | - | chr10 | 88689162 | - |
| TMEM52 | chr1 | 82897536 | + | chr1 | 82897534 | + |
| TP53 | chr5 | 42938944 | - | chr5 | 42938943 | - |
| TPM3 | chr7 | 3488208 | - | chr7 | 3488207 | - |
| XPO1 | chr10 | 33172062 | - | chr10 | 33172056 | - |

**Supplementary Table 2. RetroCNVs identified by sideRETRO in individuals from the 1000 Genomes Project.** Total retroCNVs identified per individual using whole-genome sequencing (WGS) and whole-exome sequencing (WXS) data. RetroCNVs with SR>2 (WGS) and DP>30 (WXS) were used.

| CHROM | POS | TYPE | INFO |
|---|---|---|---|
| chr1 | 9061389 | WGS | SVTYPE=INS;PG=BTBD7;PGTYPE=1;INTRONIC=SLC2A5;DP=91;SR=27 |
| chr1 | 221182809 | WGS | SVTYPE=INS;ORHO=0.348324;POLARITY=+;PG=ANO5;PGTYPE=1;DP=72;SR=64 |
| chr11 | 38791107 | WGS | SVTYPE=INS;ORHO=0.790591;POLARITY=-;PG=PCMTD1;PGTYPE=1;DP=370;SR=43 |
| chr11 | 58199876 | WGS | SVTYPE=INS;PG=TENM4;PGTYPE=1;NEAR=OR9Q2;DP=66;SR=13 |
| chr11 | 108715020 | WGS | SVTYPE=INS;ORHO=0.517508;POLARITY=-;PG=SKA3;PGTYPE=1;INTRONIC=DDX10;DP=134;SR=10 |
| chr12 | 45729702 | WGS | SVTYPE=INS;PG=ABCC1;PGTYPE=1;DP=13;SR=13 |
| chr12 | 125316601 | WGS | SVTYPE=INS;ORHO=-0.794070;POLARITY=-;PG=TDG;PGTYPE=1;DP=210;SR=36 |
| chr13 | 43495690 | WGS | SVTYPE=INS;ORHO=-0.572773;POLARITY=-;PG=TYRO3;PGTYPE=1;INTRONIC=ENOX1;DP=30;SR=3 |
| chr13 | 105145989 | WGS | SVTYPE=INS;PG=ANPEP;PGTYPE=1;DP=94;SR=27 |
| chr15 | 39702423 | WGS | SVTYPE=INS;PG=BAZ2A/RBMS2;PGTYPE=2;INTRONIC=FSIP1;DP=339;SR=28 |
| chr15 | 40561980 | WGS | SVTYPE=INS;ORHO=-0.830354;POLARITY=-;PG=CBX3;PGTYPE=1;INTRONIC=CCDC32;DP=204;SR=32 |
| chr16 | 8648136 | WGS | SVTYPE=INS;PG=SRGAP1;PGTYPE=1;EXONIC=METTL22;DP=28;SR=26 |
| chr2 | 3884044 | WGS | SVTYPE=INS;PG=ZNF664/ZNF664;PGTYPE=2;DP=50;SR=3 |
| chr2 | 54627029 | WGS | SVTYPE=INS;ORHO=0.552852;POLARITY=+;PG=CYB5D2;PGTYPE=1;INTRONIC=SPTBN1;DP=26;SR=3 |
| chr2 | 115619091 | WGS | SVTYPE=INS;ORHO=-0.785717;POLARITY=-;PG=SET;PGTYPE=1;INTRONIC=DPP10;DP=27;SR=5 |
| chr5 | 180209151 | WGS | SVTYPE=INS;PG=STK39;PGTYPE=1;EXONIC=RASGEF1C;DP=11;SR=11 |
| chr6 | 13925069 | WGS | SVTYPE=INS;PG=EXOC3;PGTYPE=1;INTRONIC=RNF182;DP=114;SR=8 |
| chr6 | 160219053 | WGS | SVTYPE=INS;PG=FRMPD3;PGTYPE=1;INTRONIC=SLC22A2;DP=12;SR=3 |
| chr7 | 49592789 | WGS | SVTYPE=INS;ORHO=0.537378;POLARITY=+;PG=WWP1;PGTYPE=1;DP=31;SR=7 |
| chr9 | 33130550 | WGS | SVTYPE=INS;PG=MRPS18A;PGTYPE=1;INTRONIC=B4GALT1;DP=40;SR=7 |
| chr1 | 9061389 | WXS | SVTYPE=INS;ORHO=-0.423754;POLARITY=+;PG=BTBD7;PGTYPE=1;INTRONIC=SLC2A5;DP=352;SR=51 |
| chr1 | 221182809 | WXS | SVTYPE=INS;ORHO=0.384514;POLARITY=+;PG=ANO5;PGTYPE=1;DP=33;SR=31 |
| chr11 | 108715020 | WXS | SVTYPE=INS;PG=SKA3;PGTYPE=1;INTRONIC=DDX10;DP=35;SR=4 |
| chr15 | 40562218 | WXS | SVTYPE=INS;IMPRECISE;CIPOS=-238,238;PG=CBX3;PGTYPE=1;INTRONIC=CCDC32;DP=56 |
| chr7 | 137300977 | WXS | SVTYPE=INS;PG=HSPE1;PGTYPE=2;INTRONIC=PTN;DP=33;SR=1 |
| chr9 | 33130550 | WXS | SVTYPE=INS;PG=MRPS18A;PGTYPE=1;INTRONIC=B4GALT1;DP=157;SR=15 |
